# Diversity in the Utilization of Different Molecular Classes of Dissolved Organic Matter by Heterotrophic Marine Bacteria

**DOI:** 10.1101/2024.02.13.580157

**Authors:** Shira Givati, Elena Forchielli, Dikla Aharonovich, Noga Barak, Osnat Weissberg, Natalia Belkin, Eyal Rahav, Daniel Segrè, Daniel Sher

## Abstract

Heterotrophic marine bacteria utilize and recycle dissolved organic matter (DOM), impacting biogeochemical cycles. It is currently unclear to what extent distinct DOM components can be utilized by different heterotrophic clades. Here, we ask how a natural microbial community from the Eastern Mediterranean Sea responds to different molecular classes of DOM. These molecular classes - peptides, amino acids, amino sugars, disaccharides, monosaccharides and organic acids - together comprise much of the biomass of living organisms, released upon their death as DOM. Bulk bacterial activity increased after 24-hours for all treatments relative to the control, while glucose and ATP uptake decreased or remained unchanged. The relative abundance of several bacterial families, assessed using 16S rRNA amplicon sequencing, increased in some treatments: peptides promoted an increase in *Pseudoalteromonadaceae*, disaccharides promoted both *Pseudoalteromonadaceae* and *Alteromonadaceae*, and most other treatments were dominated by *Vibrionaceae*. While some results were consistent with recent laboratory-based studies, for example *Pseudoalteromonadaceae* favoring peptides, other clades behaved differently. *Alteromonadaceae*, for example, grew well in the lab on many substrates but dominated in seawater samples when disaccharides were added. These results highlight the diversity in DOM utilization among heterotrophic bacteria and complexities in the response of natural communities.

**Importance:** The marine DOM pool contains numerous molecular classes, which change depending on the phytoplankton species, environmental conditions and interactions with other microbes, viruses and predators. In turn, the availability of these macromolecular pools affects the composition and function of the whole microbial community. Tracing the path between different carbon sources to specific microbes is another step towards revealing the dynamic interaction between bacteria and the DOM pool. This is especially important in warm and oligotrophic marine systems (e.g., Eastern Mediterranean Sea) where nutrients are scarce and may therefore affect microbial activity and growth.

## Introduction

Life depend on organic carbon, and the entire Earth system relies on the flow of carbon from abiotic to biotic reservoirs, conversion into organic compounds, and subsequent remineralization back to carbon dioxide (Schlesinger & Bernhardt, 2013; Summons, 1993). In the ocean, the carbon cycle begins when autotrophic microorganisms convert carbon dioxide or bicarbonate dissolved in seawater into organic carbon molecules. The organic carbon is then respired to meet the organisms’ energy needs or incorporated into biomass, mostly as macromolecular pools. In phytoplankton, between ∼30-65% of dry weight is composed of protein, up to 20% is nucleic acids, and up to ∼45% is in the form of various carbohydrates (Vargas *et al*., 1998; Geider and La Roche, 2002; Finkel *et al*., 2016). These macromolecules are expected to be released into the environment when cells die from viral lysis (Weinbauer, 2004; Kuhlisch *et al*., 2021; Moran *et al*., 2022) or predation (“sloppy feeding”, (Møller et al., 2003; Møller, 2007)). They are also released when the cells are alive, for example due to exudation of dissolved organic matter (DOM) (Thornton, 2014; Lopez *et al*., 2016; Roth-Rosenberg *et al*., 2021), or release of vesicles (Biller *et al*., 2014).

Overall, the DOM pool has been estimated to contain tens of thousands of compounds (Hertkorn *et al*., 2013). Heterotrophic microbes in the ocean depend, to a large extent, on these different organic compounds as a principal source of nutrients and energy (Larsson and Hagström, 1979), and the composition of DOM was shown to influence microbial community function (Pinhassi *et al*., 2004; Gómez-Consarnau *et al*., 2012; Becker *et al*., 2014; Bryson *et al*., 2017; Pontiller *et al*., 2020). The exact mechanisms of this relationship are not well established, but the partitioning of organic carbon fractions among various heterotrophs is thought to play a significant role in the resulting community structure (Sarmento and Gasol, 2012; Bryson *et al*., 2017; Pontiller *et al*., 2020; Ferrer-González *et al*., 2021). Heterotrophs may be specialized for specific molecular classes of the DOM (herein referred to as molecular classes) pool or degrade them with different efficiencies (Gómez-Consarnau *et al*., 2012; Sarmento *et al*., 2016). Organic molecules also serve as important signaling currencies (Keller and Surette, 2006) and changes in the chemical environment alter microbial interactions (Cude et al., 2012; D’Souza et al., 2018; Dittmar & Arnosti, 2018), adding a layer of complexity to the already complex heterotroph-DOM relationship. One example is microbial cross-feeding through syntrophic interaction, that can alter both DOM and microbial composition (Morris *et al*., 2013).

Tracing the path of specific molecular classes between microbes would aid our understanding of community function, but efforts in this area are hindered by the incredible complexity of marine organic matter (Moran *et al*., 2016; Kharbush *et al*., 2020). Previous studies analyzing organic matter degradation in environmental samples have either utilized algal exudates (e.g. (Sarmento and Gasol, 2012; Sarmento *et al*., 2016; Eigemann *et al*., 2022)), or used specific molecular classes such as amino acids (Keil and Kirchman, 1991; Middelboe *et al*., 1995; Church *et al*., 2000; Mary *et al*., 2008; Zubkov *et al*., 2008), DMSP (Ruiz-González *et al*., 2012), glucose (Rich *et al*., 1996; Church *et al*., 2000; Eilers *et al*., 2000; Kirchman *et al*., 2000; Haider *et al*., 2023), pyruvate and acetate (Baltar *et al*., 2016), and phosphonates (Dyhrman *et al*., 2006; Feingersch *et al*., 2012; Sosa *et al*., 2019). Several studies also compared the effect of adding multiple molecular classes to natural communities, revealing differential assimilation of substrates between taxa (Bryson *et al*., 2017), and substantial transcriptional responses, which differed between molecular classes along with taxon-specific responses (Pontiller *et al*., 2020). Thus, despite significant work on the role of the specific molecules mentioned above, much less is known about other, broadly defined, molecular classes such as different types of carbohydrates, peptides etc. This is important, as the dynamics of heterotrophic bacteria are often expected to be controlled by the availability of these pools (Teeling *et al*., 2012).

Here, we examine if and how natural microbial communities from the ultra-oligotrophic Eastern Mediterranean Sea (EMS) respond to addition of different molecular classes: peptides, amino acids, amino sugars, disaccharides, monosaccharides, and organic acids. To this end, we characterized changes in community composition (16S rRNA expression) and activity (different enzymatic assays) in response to amendments of different molecular classes. We also compared our results with a recent study that has shown that marine heterotrophic bacteria from a diverse collection, grown in lab batch culture, respond in different ways to the same molecular classes (Forchielli et al., 2022). While these responses could be broadly clustered by taxonomy, the molecular classes ‘preferences’ were more related to the presence of specific metabolic pathways (Forchielli et al., 2022). This allowed us to ask whether similar preferential utilization of molecular classes could be observed in a system that captures the diversity and complexity of natural microbial ecosystem.

## Materials & Methods

### Overview and experimental design

Surface seawater (from 10m depth) were collected at the continental slope of the EMS, where the bottom depth was ∼800m (Latitude = 32 30.26 N; Longitude = 034 37.52 E) during November 11, 2019. Seawater were amended in the lab, approximately ∼15 hours after collected onboard. Seawater were amended with six defined molecular classes of DOM (Table 1), each at a concentration of 25 µM (media composition and actual concentrations added to the bottles are listed in Supplementary Table 2). In addition, inorganic nutrients were also added to make sure that the heterotrophic bacteria were not N or P limited and could thus utilize the organic molecular classes as carbon sources. Thus, ∼130 µM of NH_4_ and 5 µM of PO_4_ were added, resulting in N:P ratio of ∼27:1, in accordance with the seep water masses of the EMS (Ben Ezra *et al*., 2021) (Table 1). Seawater were incubated in quadruplicate in 4.5L bottles in running seawater pools to maintain ambient temperature of ∼25°C in the dark for 24 hours. Samples were taken at both T_0_ and T_24_ from each incubation bottle for: microbial cell abundance using flow cytometry, bacterial productivity using radio labeled leucine, glucose and ATP uptake using radioisotopes, and alkaline phosphatase activity (for discussion on the differences between T_0_ and T_24_ see supplementary text 1). In addition, RNA samples for 16S amplicon sequencing were taken at T_24_.

**Table 1.** Summary of the different experimental treatments, including net added concentrations and elemental ratios, performed on Nov. 2019 at the surface water of the EMS.

| | C [μM] | NH<sub>4</sub> [μM] | PO<sub>4</sub> [μM] | C:N | C:P | N:P |
| --- | --- | --- | --- | --- | --- | --- |
| **Control** | 0 | 0 | 0 | --- | --- | --- |
| **Inorganic N+P** | 0 | 133 | 5 | --- | --- | 27 |
| **Peptides** | 25 | 133 | 5 | 0.19 | 5 | 27 |
| **Amino Acids** | 25 | 141 | 5 | 0.18 | 5 | 28 |
| **Disaccharides** | 25 | 133 | 5 | 0.19 | 5 | 27 |
| **Monosaccharides** | 25 | 133 | 5 | 0.19 | 5 | 27 |
| **Amino sugars** | 25 | 137 | 5 | 0.18 | 5 | 27 |
| **Organic Acids** | 25 | 133 | 5 | 0.19 | 5 | 27 |

### Bacterial productivity

Bacterial productivity (BP) was estimated using the ^3^H-leucine incorporation method (Simon and Azam, 1989) and is described in detailed by (Reich *et al*., 2022). Triplicate 1.7ml of seawater were incubated with ^3^H-leucine (20 nmol leucine L^-1^) for 4h at ambient temperature of ∼25°C in the dark immediately after sampling. The incorporation was terminated by adding 100 μl of cold trichloroacetic acid (TCA). Killed blanks containing seawater, the radiolabeled leucine and TCA added immediately upon collection (no incubation) were also undertaken and subtracted from the sample reads. The samples processed using the micro-centrifugation protocol (Simon and Azam, 1989). Scintillation cocktail with high affinity to beta radiation (ULTIMA-GOLD) was added before counted using TRI-CARB 2100 TR (PACKARD) scintillation counter.

### Glucose and ATP bulk uptake

Bulk uptake rates of Adenosine 5’-triphosphate and Glucose D[-^3^H(N)] were estimated using modified assay adapted from (Alonso-Sáez *et al*., 2012). Triplicate 20 ml of seawater were spiked immediately after sampling with either ^3^H-Adenosine Triphosphate or ^3^H-D-Glucose (added concentration of 0.2 nM for each substrate, 0.1548 μCi and 0.1836 μCi in sample respectively) and incubated for 4h at ambient temperature in the dark. The incorporation was terminated by filtration of the samples onto 0.22 μm polycarbonate filters followed by subsequent rinsing with 5 ml of 0.22 μm filtered seawater. Killed blanks containing seawater, the radiolabeled substrate and formaldehyde (2% final concentration) added immediately upon collection and 15 minutes prior to the spike were also undertaken and subtracted from the sample reads. A scintillation cocktail with high affinity to beta radiation (ULTIMA-GOLD) was added before counted using TRI-CARB 2100 TR (PACKARD) scintillation counter.

### Alkaline phosphatase activity (APA)

APA was determined by the 4-methylumbeliferyl phosphate (MUF-P: Sigma M8168) method (Thingstad and Mantoura, 2005). After the addition of substrate to a final concentration of 50 µM, samples were incubated in the dark at ambient temperature for 3/4 hour.

### RNA extraction, DNA digestion, cDNA synthesis

Bacterial community/activity was assessed by 16S rRNA. Approximately ∼4.4 L of SW from each incubation bottle was filtered directly onto 0.2 μm filters, added with RNAsave and stored in −80℃ until extraction. RNA was extracted using RNeasy® PowerWater® Kit following the manufacture protocol, including DNase treatment. RNA was transcribed into cDNA using iScript cDNA Synthesis.

### 16S rRNA sequencing

Sequencing was performed using primers 515F and 926R (Walters *et al*., 2015). Amplicons were generated using a two-stage polymerase chain reaction (PCR) amplification protocol. First stage PCR amplification was carried out in 25 μl reactions using MyTaq Red Mix (BIO-25044, Meridian Bioscience). The amplification parameters were set as follows: 95° C for 5 min, followed by 28 cycles at 95°C for 30s, 50°C for 30s, and 72°C for 1 min. A final, 5-min elongation step was performed at 72°C. Products were verified on a 1% agarose gel before moving forward to the 2nd stage. One microliter of PCR product from the first stage amplification was used as template for the 2nd stage, without cleanup. Cycling conditions were 98°C for 2 minutes, followed by 8 cycles of 98°C for 10s, 60°C for 1min and 68°C for 1min. Libraries were then pooled and sequenced with a 15% phiX spike-in on an Illumina MiSeq sequencer employing V3 chemistry (2×300 base paired-end reads). Library preparation and sequencing were performed at the Genomics and Microbiome Core Facility (GMCF; Rush University, IL, USA).

### 16S rRNA sequence analysis

All sequences were analyzed using Dada2 pipeline (Callahan *et al*., 2016) and the software packages R (R Development Core Team, 2011) and Rstudio (R Team, 2020). Forward and backward primers were trimmed, forward and backward reads truncated after quality inspections to 250 and 230 bases respectively. After sequences merging, a consensus length only between 400 and 430 bases was accepted. Finally, ASVs that have less than 100 in total (all samples) were removed. Silva database version 138 (Quast *et al*., 2012) was used for taxonomic assignment. All chloroplasts, mitochondria, archaea, eukaryotes and Amplicon sequence variants (ASVs) without any taxonomic affiliation were discarded from downstream analyses.

### Statistics

Data pre-processing and statistical analyses were performed using the R statistical programming language (R Development Core Team, 2011; R Team, 2020). One-way ANOVA and post-hoc Tukey test were performed to compare between the treatments on each time points separately (Figures 2 and 3, Supplementary Table 1) using multcompView (Graves et al., 2019) and dplyr (Wickham et al., 2023) packages. Paired t-test with Bonferroni correction was used to compare the means between time points of each treatment (Supplementary Figures 3 and 4) using the packages tidyverse (v1.3.0; (Wickham et al., 2019), rstatix (Kassambara, 2023b) and ggpubr (Kassambara, 2023a). PERMANOVA test was used to test the significance of the NMDS plots distribution (Figure 4). Mantel test with 9999 permutations examining the Spearman’s correlation was used to test the correlation between the BioCyc and 16S results (Figure 6) using vegan package (Oksanen et al., 2022). Plots were originated using ggplot2 package (Wickham, 2016).

## Results

### Initial seawater characteristics and experimental setup

The marine microbial community to which we added different molecular classes was from a transition region between coastal and open-ocean, oligotrophic waters in the EMS (Figure 1A). The experiment was performed using surface water (10m depth) collected from an offshore location (∼800m water depth). Water was collected at the end of fall, moving into winter, after 11-12 days of constant Easterly or Southerly winds and no rain. The vertical temperature profile indicates that the water column was still stratified, with ∼25°C in the upper 50m (Figure 1C) and salinity of ∼39.5 ppt throughout (Supplementary Figure 1B). Orthophosphate in the surface water was below the limit of detection (Figure 1D), and alkaline phosphatase activity (APA) was relatively high, suggesting phosphorus-limitation for microbes (Figure 1E). Surface NO_3_ + NO_2_ concentration was 0.19µM (Figure 1D), and surface primary productivity values were 0.35±0.03 µg C L^-1^ h^-1^ (Supplementary Figure 2E), both of which were somewhat higher than previously reported for a parallel season in the open-ocean Eastern Mediterranean (Ben Ezra *et al*., 2021; Reich *et al*., 2022). This suggests that despite the stratification, winter mixing had begun, in agreement with results from a time-series study from a nearby location (Ben Ezra *et al*., 2021).

**Figure 1.**
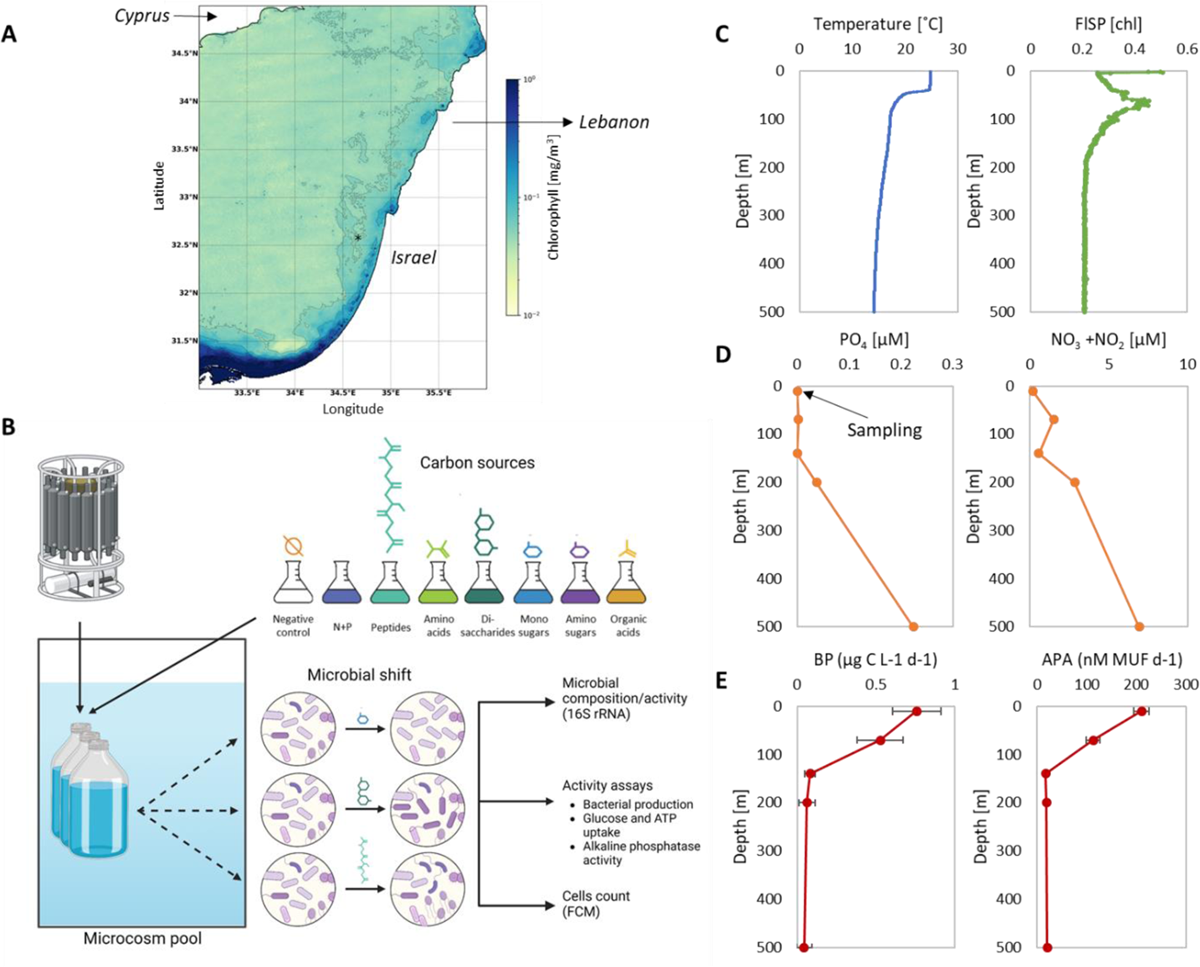
Oceanographic context and experimental setup. (A) Chlorophyll satellite image of the location site (marked with asterisk) in the EMS from which surface water were collected for the microcosm experiment (32 30.26 N; 034 37.52 E). (B) Overview of the experimental setup. (C-E) Depth profiles at the location site. (C) Temperature and chlorophyll (from CTD). (D) PO_4_ and NOx (NO_3_ + NO_2_) (Ben Ezra *et al*., 2021). (E) Bacterial productivity (BP) and alkaline phosphatase activity (APA). Seawater samples were taken from 10m depth.

**Figure 2.**
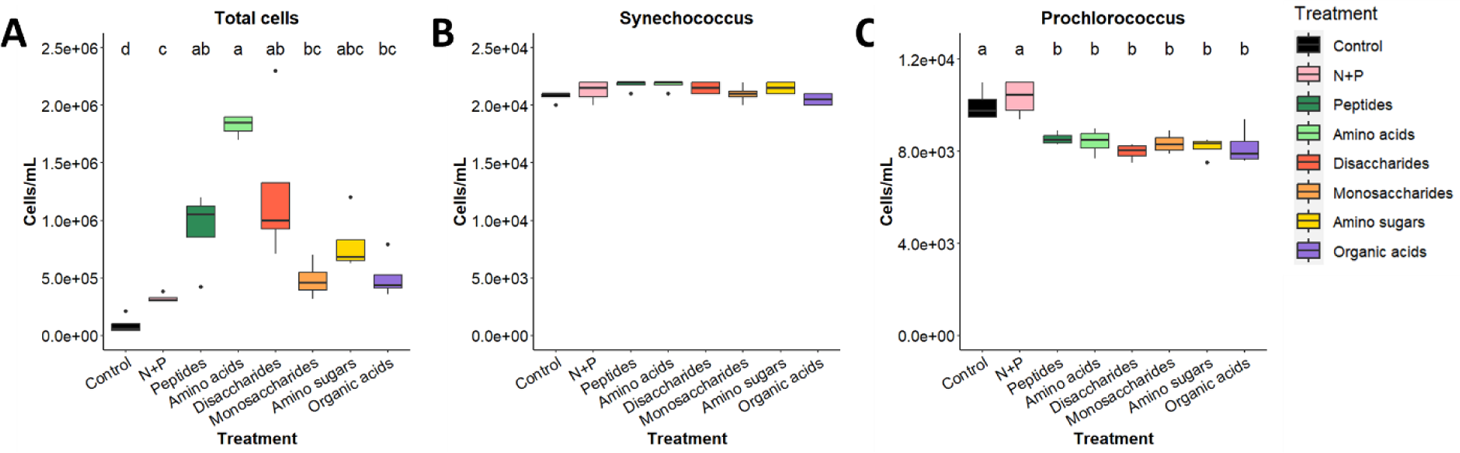
Changes in microcosm bacterial abundance at T_24_. (A). Total bacterial abundance. (B) *Synechococcus*. (C) *Prochlorococcus*. Box-Whisker plots show the interquartile range (25th–75th percentile) of the data set. The horizontal line within the box represents the median value (N=4). The different letters above the box plots indicate statistically significant differences among the treatments (one-way ANOVA and post-hoc Tukey test, *p* < 0.05). There was no significant difference between the treatments for *Synechococcus*.

**Figure 3.**
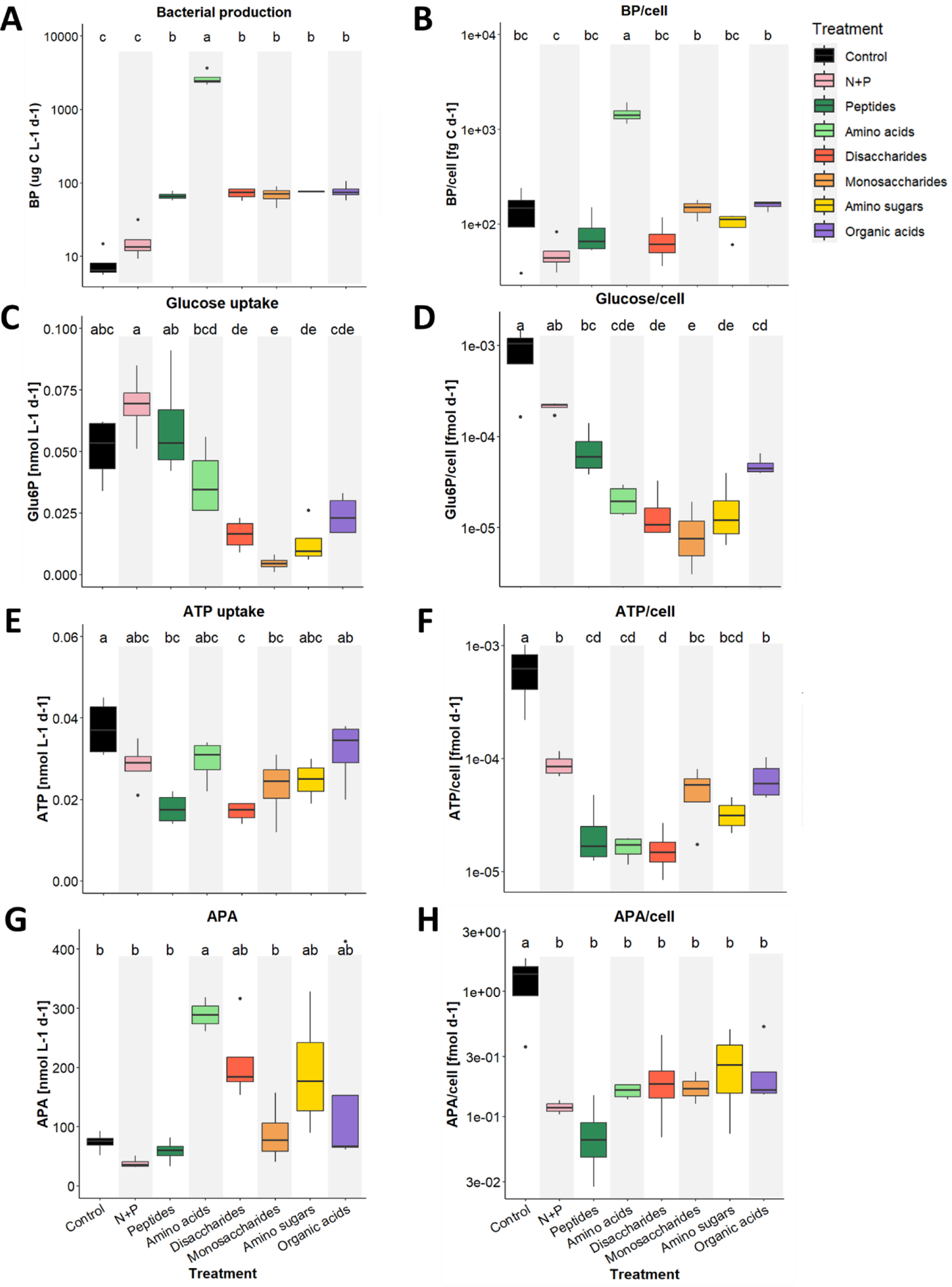
Different parameters of bacterial activity as bulk and per-cell at T_24_. (A, B) Bacterial production. (C, D) Glucose uptake. (E, F) ATP uptake. (G, H) Alkaline phosphatase activity (APA). Note the logarithmic Y-axis on A and on the right panel. Box-Whisker plots show the interquartile range (25th–75th percentile) of the data set. The horizontal line within the box represents the median value (N=4). The different letters above the box plots indicate statistically significant differences among the treatments (one-way ANOVA and post-hoc Tukey test, *p* < 0.05). Bulk and per-cell BP uptake rates with amino acids were corrected to consider the “cold” Leucin concentration (1.9 µM, see supplementary text 2).

**Figure 4.**
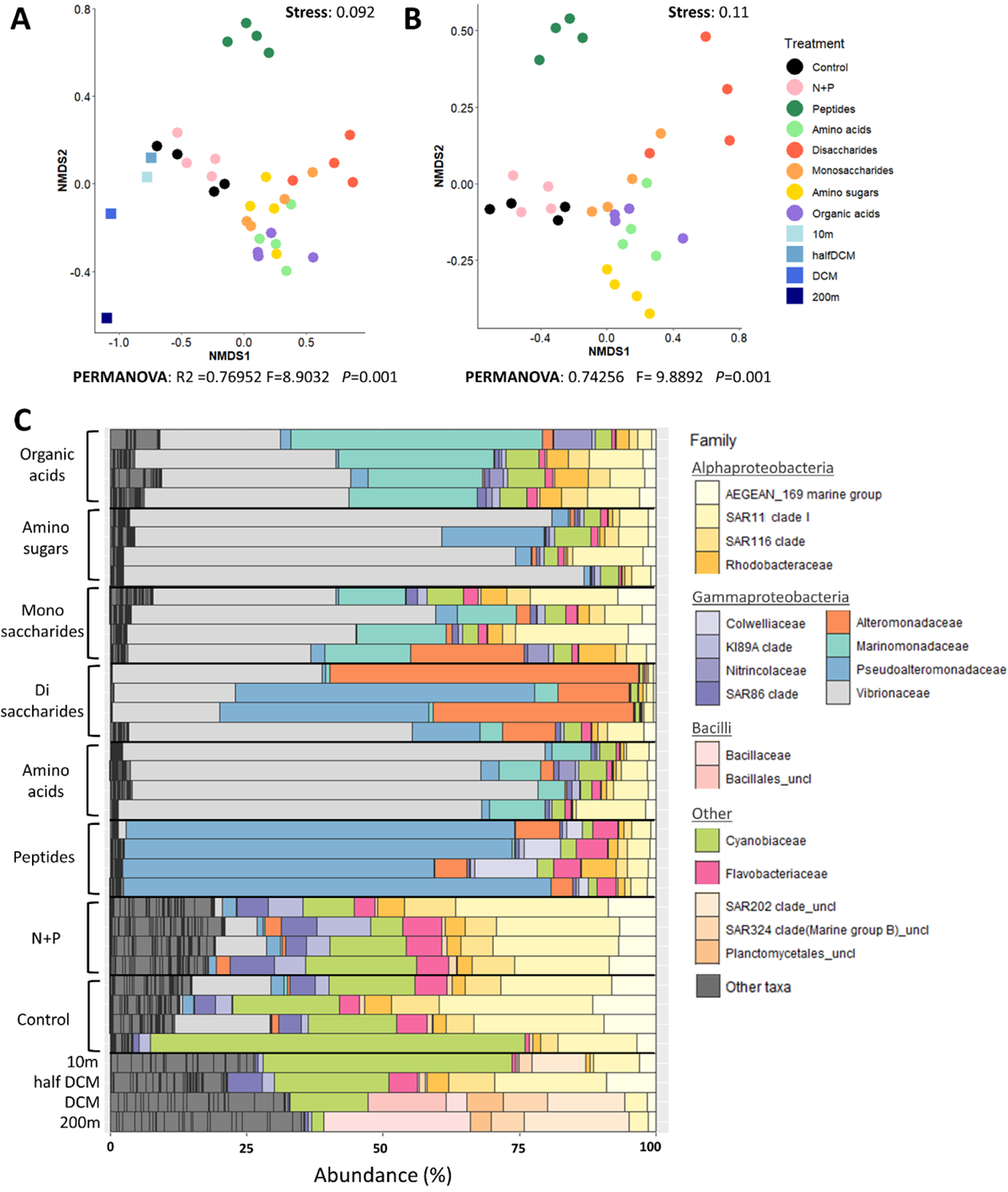
Results of 16S rRNA analysis. Nonmetric multidimensional scaling (NMDS) plot, based on Bray-Curtis dissimilarity for 16S amplicon for (A) all microcosm and cruise (field) samples and (B) microcosm samples. Only ASVs that were on average greater than 1% of the community are considered. Results of PERMANOVA test are shown in both panels. (C) Stacked bar chart showing relative proportions (percentages of 16S rRNA sequences) of microbial community at T24 at the family level. *Pseudoalteromonadaceae* responded primarily to peptides, and (with *Alteromonadaceae*) to disaccharides; *Marinomonadaceae* responded to organic acids, and *Vibrionaceae* to everything but peptides. Uncl=uncultured. Other taxa=families not greater than 5% in any sample.

To characterize the response of this natural community to different classes of labile DOM, we amended the collected seawater with six different molecular classes: peptides, amino acids, monosaccharides, disaccharides, amino sugars and organic acids, each at 25μM (see Supplementary Table 2 for detailed media composition, which mirrored those of a previous in vitro experiment (Forchielli et al., 2022)). We also included a negative control without any modification, and a control amended only with NH_4_ and PO_4_, which were added to mitigate any potential inorganic nutrient limitation (Figure 1B, Table 1).

### Microbial Abundance

After 24 hours incubation, total bacterial abundance was ∼5×10^4^ cells ml^-1^ in the un-amended controls, and significantly increased (∼3.5 fold) in the N+P addition (Figure 2A). Peptides, amino acids and disaccharides all elicited an additional ∼3-5.5 fold increase above the N+P treatment bottles (*P*-value<0.05, one-way ANOVA), whereas monosaccharides, amino sugars and organic acids elicited smaller increases which were not statistically significant.

Two autotrophic prokaryotes groups were identified: *Synechococcus* ranging from 2-2.5×10^5^ cells ml^-1^ (Figure 2B) and *Prochlorococcus* ranging from 7.5×10^3^ - 1.3×10^4^ cells ml^-1^ (Figure 2C). The addition of molecular classes reduced *Prochlorococcus (*but not *Synechococcus*) counts relative to the un-amended and N+P controls (Figures 2B and C).

### Bacterial activity assays

BP, a commonly used general measure of heterotrophic bacterial activity, is measured by the incorporation of radiolabeled leucine, and thus represents the uptake of amino acids and their incorporation into biomass. After 24 hours bulk BP significantly increased for all molecular classes compared to both the un-amended and N+P controls (Figure 3A), with the largest increase observed with amino acids, as previously shown (Middelboe *et al*., 1995; Zubkov *et al*., 2008). In contrast to the bulk BP, BP/cell increased significantly in relation to the un-amended and N+P controls only after the addition of amino acids (Figure 3B).

As opposed to bacterial productivity, bulk uptake rate of glucose and ATP either decreased or did not change relative to the control and N+P (Figure 3 C, E), and the per-cell values significantly decreased (Figure 3 D, F). Notably, already at T_0_ (right after the addition of the macromolecular classes) glucose uptake rates for monosaccharides, disaccharides and amino sugars were significantly lower compared with the other treatments (*P*-value<0.05, one-way ANOVA, supplementary Table 1, Supplementary Figure 4C). Since glucose was one of the monosaccharides used for these experiments, it is possible that the decrease in the uptake of the radiolabeled sugar is due to competition between the “hot” (radiolabeled) and “cold” substrate. However, the decrease in disaccharides and amino sugars is less expected and may be due to cross-reactivity of the transporters or to catabolite repression, as discussed in more detail in supplementary text 2.

In addition to measuring the uptake rates of amino acids, glucose and ATP, we also measured extracellular alkaline phosphatase activity (APA). Higher AP activity is expected when PO_4_ concentration are low (Cembella et al., 1982), and thus a good estimator for phosphorous starvation. Initial APA values were similar across all treatments, but lower than measured *in-situ* at the sampling site (∼60 compared to ∼240 nM MUF/day in the surface water, compare Figure 3G and Figure 1E). Bulk APA significantly increased compared to the un-amended and N+P controls only for amino acids, but some degree of increase was also observed for most other treatments (Figure 3G), possibly as a result of the increased bacterial abundance. In contrast, APA/cell significantly decreased in all treatments compared to the control, which is expected since phosphate was added (Figure 3H).

### Changes in microbial community composition

To determine whether the microbial community composition was altered in response to the different added molecular classes, we amplified and sequenced the 16S rRNA (i.e. from extracted total RNA). Observed changes are therefore due to both changes in ribosomal RNA gene expression (associated with increases in activity or growth rate) and changes in cell numbers (Salazar *et al*., 2019). The un-amended and N+P controls samples grouped together and close to the surface (10m) and half-DCM (70m) samples collected in the field (Figure 4A). The field, control and N+P samples were also more diverse than the samples to which molecular classes were added (alpha diversity, Supplementary Figure 5), and dominated by cyanobacteria (mostly *Synechococcus* but also *Prochlorococcus*) and alphaproteobacteria (SAR11, SAR 116 or AEGEAN-169) (Figure 4C). The 10m sample, from which water were taken for the experiment, was composed of ∼45% cyanobacteria, ∼10% SAR202 and ∼8% SAR11 (clade 1), with other families below 5%. In addition to cyanobacteria and alphaproteobacteria, the un-amended and N+P controls also had more Flavobacteria (∼6%) and few gammaproteobacteria families: *Vibrionaceae* (∼5-15%), and (for the N+P control) KI89A and SAR86 clades (∼5-15%).

The addition of each molecular class elicited a different change in the microbial community rRNA profile, and in many of these cases it resulted in the growth of a different clade of heterotrophic bacteria (Figure 4 B, C). The peptide treatment, which was dominated by *Pseudoalteromonadacea,* clearly differed from all others in the NMDS ordination. Two other treatments, disaccharides and amino sugars also each group separately. The disaccharide treatments were dominated by *Alteromonadaceae*, *Pseudoalteromonadaceae* and to some extent *Vibrionaceae*, while the amino sugars treatment was dominated by *Vibrionaceae*. The community growing on organic acids was also dominated by *Vibrionaceae* but also by *Marinomonadaceae*, which were also quite abundant in the amino acids and most monosaccharides treatments. Overall, in terms of relative abundance, the dominant family in all treatment except peptides was *Vibrionaceae*, with the highest percentages observed for amino acids and amino sugars (∼75%), followed by monosaccharides, disaccharides and organic acids (∼25-50%) (Figure 4C).

Within some of the families there were also higher-resolution patterns in the relative abundance of specific Amplicon Sequence Variants (ASVs). The most abundant family, *Vibrionaceae,* was comprised of ∼6-8 prevalent ASVs, three of which were specific for *Photobacterium sp*. (a common fish pathogen), and the others could not be identified to a higher resolution than the family level. The *Photobacterium* ASVs were relatively abundant only in response to amino sugars, and actually decreased in relative abundance in other treatments, whereas the other ASVs were unspecific and found among all treatments without any clear patterns (Supplementary Figures 6, 7A). In contrast, *Pseudoalteromonadaceae* had 3 main ASVs with ∼90-97% identity. The first and unspecific ASV, ASV3, was very dominant for peptides while the other two, one of which matches the genus *Psychrosphaera*, were mostly found in two of the disaccharide’s treatment (Supplementary Figure 7B). In contrast, no clear intra-family patterns were observed for *Alteromonadaceae* and *Marinomonadaceae* (Supplementary Figure 7C, D). ASV belonging to the four most abundant families responding to the molecular classes were not identified in the original seawater (from 10m depth), except for one ASV of *Pseudoalteromonadacea* with relative abundance of 0.03%.

### Differences between patterns of carbon source utilization in laboratory cultures and in natural communities

How do the results presented above compare with a set of laboratory experiments in which 63 selected strains of marine heterotrophic bacteria were grown with the same molecular classes (Forchielli et al., 2022)? As shown in Figure 5, the change in relative abundance of some specific families in our microcosm experiment was consistent with the growth of their cultured representatives in lab cultures. For example, *Pseudoalteromonadaceae*, the dominant family observed when peptides were added to natural seawater, also grew well on peptides in monocultures (Figure 5). *Pseudoalteromonadaceae* were also very dominant in two disaccharides samples in the microcosm, yet only one cultured strain, *Pseudoalteromonas citrea*, was able to grow on disaccharides (Forchielli et al., 2022), suggesting the potential for intra-family diversity in the ability to utilize this molecular class. *Rhodobacteraceae*, although not one of the most common families, also behaved similarly, growing well in lab monocultures on organic acids and monosaccharides and in most microcosm replicates where organic acids and monosaccharides were added (∼2-5%, compared to <2% in the other treatments). The growth of other clades, however, was less consistent between the laboratory cultures and natural communities in the microcosm. Lab cultures of *Alteromonadaceae*, for example, grew well on amino acids, disaccharides and amino sugars, whereas they dominated in the microcosms only when disaccharides were added. In addition, *Vibrionaceae,* which were common in all microcosm treatments except peptides, grew in lab cultures mostly on amino acids and peptides (Figure 5, (Forchielli et al., 2022)).

**Figure 5.**
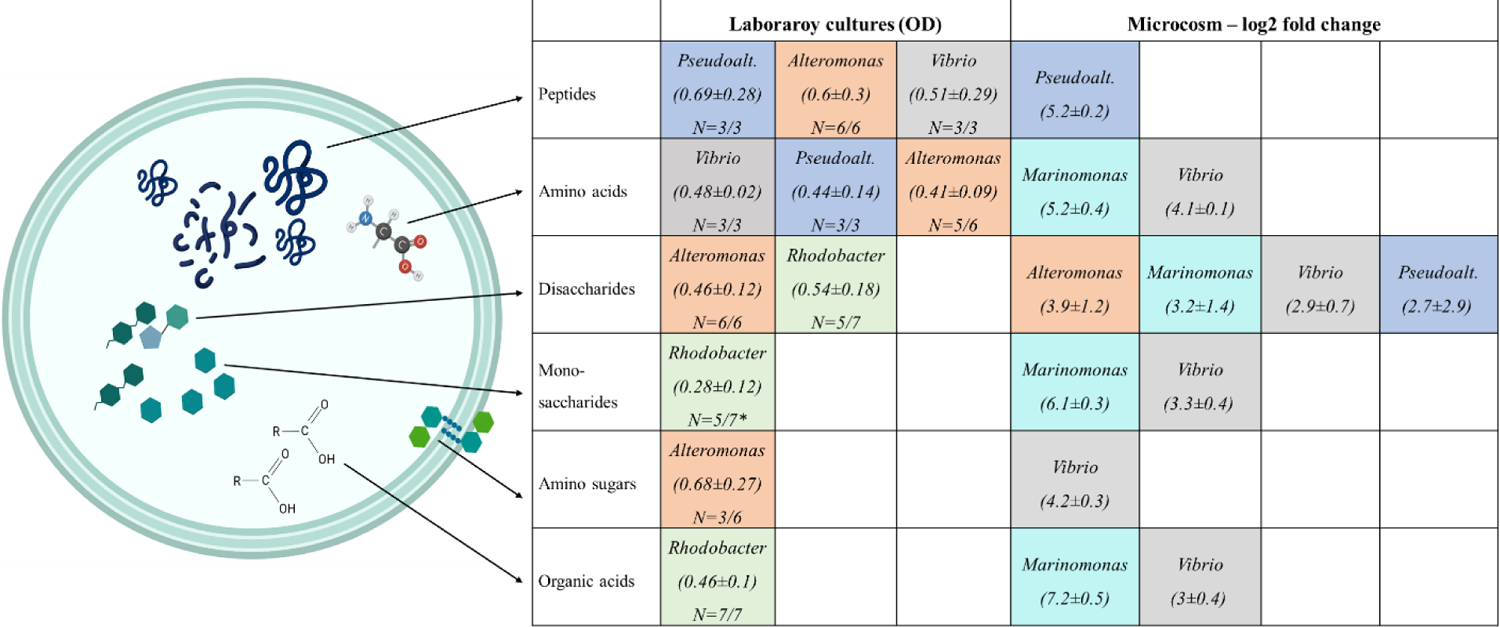
Bacterial growth on phytoplankton-derived molecular classes in lab (monoculture) and in the microcosm (mixed community). Proteins, including **peptides** and **amino acids,** are key components in the cell and comprise up to 60% of its dry weight (Geider and La Roche, 2002). **Sugars** are energy sources, comprising in our experiments **disaccharides** and **monosaccharides**, as well as **amino sugars** such as N-acetylglucosamine (GlcNAc) which are major components of the cell wall (Konopka, 2012). **Organic acids** are common metabolites produced and consumed by different microorganisms, and genes for their utilization were shown to be upregulated in bacterioplankton following phytoplankton bloom (Rinta-Kanto *et al*., 2012). Colors in the table represent different families. For the microcosm, only families with an average (N=4) log2 fold change>1.5 are shown. In the lab, we include only genera with a minimum of 3 strains tested, and which were able to grow above average OD_600nm_ of 0.4 (Forchielli et al., 2022). N represents the number of strains able to grow on each molecular class. *Rhodobacter was the only genus to show any substantial growth on monosaccharides, and thus included in the table although its average OD was lower than 0.4.

**Figure 6.**
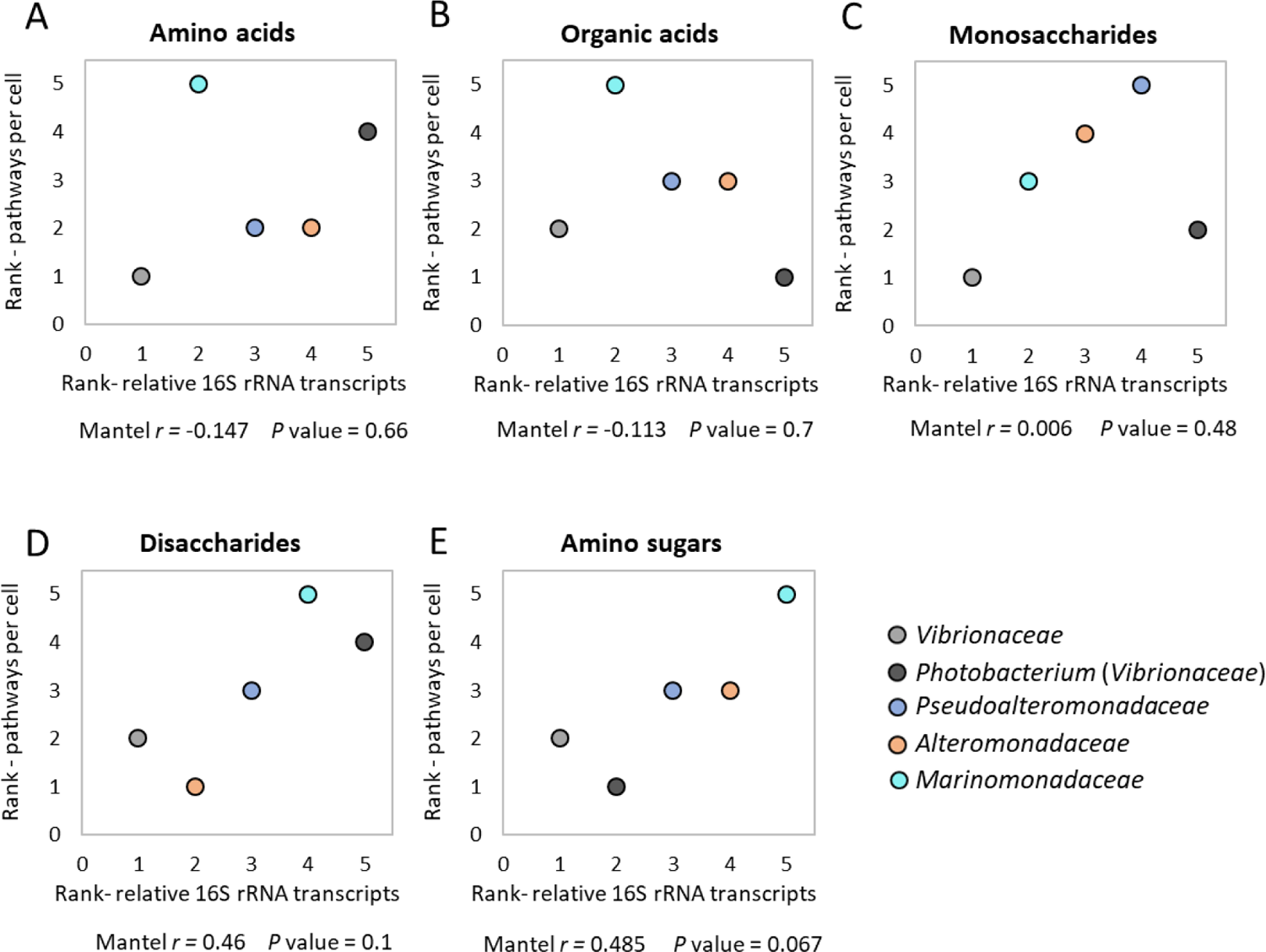
Correlation between metabolic pathways (BioCyc) and average relative abundance/activity (16S rRNA transcript amplicons) for the five most abundant families. (A) Amino acids. (B) Organic acids. (C) Monosaccharides. (D) Amino sugars (E) Disaccharides. Data shown are ranked values based on Supplementary Tables 4 and 5 and is further discussed in supplementary text 3. *r* and *P*-values are the results of Mantel test with 9999 permutations examining the Spearman’s correlation. Black lines are linear trendlines.

Is there a correlation between the number of metabolic pathways for the degradation of each carbon source type, encoded in the genome of each heterotrophic clade, and its propensity to become dominant in response to this carbon source? As shown in Figure 6, there was no statistically significant correlation between the relative increase in rRNA transcripts for each clade in response to the different molecular classes and the average number of pathways for the degradation of these molecular classes. However, disaccharides and amino sugars did show a trend for positive correlation (Figure 6D, E and supplementary text 3).

## Discussion

The goal of this study was to ask how a natural marine microbial community from oligotrophic waters responds to different molecular classes (peptides, amino acids, disaccharides, monosaccharides, amino sugars and organic acids) in terms of activity and community composition. We identified four main Gammaproteobacteria families (*Vibrionaceae, Pseudoalteromonadaceae*, *Alteromonadaceae* and *Marinomonadaceae*) that responded specifically to the addition of different molecular classes. Below we discuss these results in light of other studies asking how different types of organic matter affect marine microbial communities or individual bacterial strains, using them to illuminate how specific metabolic pathways, metabolic regulation and microbial interactions may all play a role in determining community dynamics.

### The addition of distinct molecular classes of DOM induces the growth of relatively rare but ecologically relevant heterotrophic families

The four main Gammaproteobacterial families which responded to the addition of the molecular classes are mostly rare in the original seawater community, which was dominated by alpha-proteobacteria and cyanobacteria, as previously observed (Haber et al., 2022; Roth Rosenberg et al., 2021). Yet these clades are sometimes relatively abundant in specific oceanic niches such as marine particles (Haber et al., 2022; Lyons et al., 2007; Roth Rosenberg et al., 2021; Takemura et al., 2014). They also tend to become dominant during mesocosm experiments in response to additions of both specific molecules such as glucose (Eilers *et al*., 2000; Haider *et al*., 2023) and complex mixtures (e.g. high molecular weight DOM, (Sosa *et al*., 2015), and see below).

Previous studies have shown that rare microorganisms have higher respiration rates compared to more abundant bacteria, suggesting that despite their overall rarity these copiotrophic bacteria have an important role in the remineralization of organic carbon (Munson-McGee *et al*., 2022).

Several previous studies have compared the responses of natural communities to the addition of different molecular classes, in relatively productive coastal or brackish locations (the California coast (Bryson *et al*., 2017), and the Baltic Sea (Pontiller *et al*., 2020). Similar to our study, different molecular classes elicited different community responses in terms of relative abundance and subsequent substrate uptake. For example, *Alteromonadaceae* responded both to polysaccharides in the California coast (Pontiller *et al*., 2020) and to disaccharides in our study. Yet in many cases the specific clades responding were different. For example, *Flavobacteria* from the California coast responded primarily to glucose and starch, and from the Baltic Sea to proteins, yet this clade was not one of the main responders in our study. It did, however, grew in the un-amended control, N+P and peptides treatment but decreased in relative abundance in the other treatments. Conversely, *Vibrionaceae*, which were the main responders to almost all molecular classes in our study, did not respond in either of the other ones. However, in another study conducted with water from the mesopelagic North Atlantic, addition of the organic acids pyruvate and acetate induced, among other, the growth of *Vibrionaceae*, as in our study (Baltar *et al*., 2016). Thus, further studies are needed in order to determine whether these differences are due to technical aspects (e.g. amount of organic matter added or incubation time), or to ecological factors such as the trophic state of the ecosystem (oligotrophic marine waters vs productive coastal and brackish) or the season sampled.

The molecular classes we chose represent a significant part of the biomass of phytoplankton, which would be released as phytoplankton die, e.g. during the late stages of a bloom. Indeed, phytoplankton blooms are often followed by a succession of heterotrophic bacteria, especially of *Roseobacters*, *Flavobacteria* and members of the Gammaproteobacteria such as *Alteromonadaceae* (Riemann *et al*., 2000; Fandino *et al*., 2001; Pinhassi *et al*., 2004; Buchan *et al*., 2014). For example, natural macroalgae blooms were shown to induce the growth of *Vibrionaceae*, possibly related to their ability to degrade brown algal polysaccharides (Takemura *et al*., 2014; Martin-Platero *et al*., 2018). In several occasions, *Vibrionaceae* were shown to comprise up to ∼50% of total microbial community (fraction of 16S amplicons, (Zhang *et al*., 2018)). *Prochlorococcus* exudates, on the other hand, dramatically induced the growth of *Pseudoalteromonadaceae* in open ocean water from the EMS (Eigemann *et al*., 2022), and based on the results presented here we speculate that this might be due to these exudates being protein- or disaccharide-rich (Figure 4). It is noteworthy also that amino acids, peptides and disaccharides induced the highest increase in cell abundance, suggesting that these molecular classes can be used to from biomass, while others can lead mainly to respiration. This could also affect the perceived changes in community composition.

### Relationship between specific metabolic pathways and microbial growth on each carbon source type

Bacterial growth depends primarily on the ability to utilize the specific nutrient sources present. We have previously shown that such metabolic preferences can be captured by the relative abundance of some metabolic pathways or a handful of key enzymes (Forchielli et al., 2022). For example, in the lab, growth on disaccharides, monosaccharides and acidic sugars was associated with an enrichment in carbohydrate metabolism pathways (e.g. galactose, starch and sucrose) and depleted in pathways for amino acid utilization. In the complex community studied here, we observed a positive (yet not significant) correlation between the genetic capacity for degrading disaccharides and amino sugars and the dominance of specific clades when these molecular classes were added (Figures 6D and E). For example, *Vibrionacea* (as well as a sub-group, *Photobacterium)* ranked first in both average amino sugars pathways per genome and in relative abundance after 24 hours (Figure 6E). This is supported to some extent by a study that tested the uptake of the amino sugar N-acetyl-D-glucosamine (NAG) among different bacteria, where no clear pattern relative to phylogeny was found, except that all 19 *Vibrionacea* took up NAG (Riemann and Azam, 2002). Similarly, both *Vibrionacea* and *Alteromonadaceae* had relatively more pathways for the utilization of disaccharides per cell, and indeed these two clades were the dominant ones in the disaccharide-treated microcosms (Figure 6D). This is partly in accordance with another incubation experiment where *Alteromonadaceae* responded to the addition of polysaccharides by growing and expressing genes related to motility and glycogen utilization (Pontiller *et al*., 2020). *Alteromonadaceae* have also been suggested to degrade complex carbohydrates in when incubated with natural organic material such as jellyfish detritus (Tinta *et al*., 2023), and several *Alteromonas* strains have been shown to degrade multiple polysaccharides in the lab (e.g. (Koch *et al*., 2019)).

In contrast to disaccharides and amino sugars, we did not observe any correlation between the relative abundance of specific clades in response to amino acids, organic acids and monosaccharides and the presence of degradation pathways for these molecular classes. When individual strains were tested in the lab (Forchielli et al., 2022), growth on these molecular classes was not associated with the number pathways for their utilization but rather with other pathways that may interact with them. For example, growth on organic acids was associated with enrichment for specific portions of the ethylmalonyl-CoA pathway, which is an alternative to the glyoxylate shunt used in growth on some organic acids (Forchielli et al., 2022). This would not have been captured in our more simple analysis.

It should be noted that we did not test here for correlations between the genetic capacity to degrade peptides and dominance when peptides were added, since growth on peptides depends primarily on their degradation by a wide range of relatively less characterized extracellular and intracellular peptidases. However, there are indications that *Pseudoalteromonas*, the dominant family growing on peptides in both laboratory monocultures and our mesocosms, is an important player in peptides degradation in marine environments (Zhao *et al*., 2012; Tang *et al*., 2020; Tinta *et al*., 2023).

### Are the effects of molecular classes on enzymatic activity and cell uptake due to changes in community structure or function?

A surprising observation in our study is that, while bulk amino acid uptake (BP) increased in all treatments, glucose and ATP uptake did not, and actually decreased on a per-cell basis (Figure 3). Similar observations were reported in seawater samples from the Gulf of Mexico, in which bulk uptake rates of glucose decreased in response to high molecular weight DOM and increased with the addition inorganic nutrients (Skoog *et al*., 1999). Glucose is considered to be a common organic molecule in the ocean (Rich *et al*., 1996), and the addition of glucose can increase heterotrophic microbial activity and viability in the EMS (Rahav *et al*., 2019). The decrease in glucose uptake in response to a wide range of molecular classes could, in principle, be explained by a shift in community composition, to one where the dominant organisms do not utilize glucose (such as some SAR11 strains, (Schwalbach *et al*., 2010)). However, since *Vibrionacea* and *Marinomonadacea* were the dominant organisms in most mesocosms, including those to which monosaccharides were added, it is less likely that the decrease in glucose uptake is due to the shift in community composition. We propose that the reduction in glucose uptake is more likely to be explained by a change in the physiology of the dominant community members, for example through catabolite repression, where the addition of one carbon source reduces the expression of pathways for the use of another (e.g. downregulation of glucose transporters).

In contrast to glucose, which cannot be used by all marine bacteria, ATP is a key metabolite in every organism, involved in thousands of metabolic reactions, and found in all cells at millimolar concentrations. ATP taken up from the environment can provide the cells with both energy and phosphorus, which is often limiting in the EMS (Ben Ezra *et al*., 2021; Reich *et al*., 2022). Similar to the decreased APA activity, the decrease in ATP uptake across all experimental conditions is most likely related to the alleviation of phosphate limitation by the addition of PO_4_ (Sebastián *et al*., 2012). This also suggests that any change in community composition likely represents bacteria able to use these the added molecular classes rather than a response to phosphorus starvation.

As opposed to glucose and ATP, the per-cell uptake rates of leucine (the amino acid used for the BP assay) remained stable after 24 hours, with the exception of the amino acid treatment, in which it increased. This could be explained by two (non-exclusive) hypotheses: (1) amino acid uptake and utilization, unlike glucose, does not undergo catabolite repression; (2) all of the (dominant) bacteria in the community can take up and utilize amino acids. In support of the second hypothesis, the genomes of more strains from the dominant bacteria in the mesocosms contain pathways for the degradation of leucine compared to glucose (67±30% and 18±30%, respectively, supplementary Table 6). Furthermore, it has been previously shown that more than ∼50% of bacterial cells take up leucin in different marine environments (Kirchman et al., 1985), compared with ∼20% taking up glucose (Alonso and Pernthaler, 2005). Regardless of whether amino acids do not undergo catabolite repression or are simply used by all the dominant bacteria, these results provide further support that amino acids are common metabolic currencies in the marine environment.

### Microbial interactions and competition- the ‘big picture’

Until now we have focused on the factors that determine whether an individual clade of bacteria responds to the addition of different carbon source types. However, in experiments with natural communities, interactions such as competition, allelopathy and syntrophy play a major role in shaping microbial composition, influencing their metabolic activity (Orphan *et al*., 2001; Morris *et al*., 2013; Corno *et al*., 2015; Datta *et al*., 2016), and thus the collective community function (Fuhrman *et al*., 2015; Graham *et al*., 2016). For example, competition for the same molecular classes and allelopathy, the process in which one organism produces compounds that influence the growth and survival of another (Long and Azam, 2001), might explain the discrepancies between the laboratory cultures (Forchielli et al., 2022) and our microcosm experiment (see further discussion in Supplementary Text 4). Indeed, all dominant families observed in our study are able to produce anti-microbial compounds (Holmström, 1999; Lucas-Elio *et al*., 2005; Jeganathan *et al*., 2013; Zoccarato *et al*., 2022), but some seem to be more efficient competitors. For example, a major inconsistency between the laboratory monocultures and mesocosms was that *Alteromonadaceae* grew well in lab mono-cultures but became dominant in microcosms only when disaccharides were added. In most other cases they were outcompeted by *Vibrionaceae*. In a systematic analysis of antagonistic interacting between marine bacteria, *Alteromonadaceae* and *Vibrionaceae* did not tent to inhibit one another (Long and Azam, 2001).

However, other studies suggest evidences for the competition between both families (Michotey *et al*., 2020), and for the advantage of *Vibrio* over *Alteromonas* when growing together on different molecular classes (Eilers *et al*., 2000). We speculate that only when *Alteromonadaceae* had a clear metabolic advantage in the disaccharides treatment it was able to successfully outcompete *Vibrionaceae*. Another case of potential allelopathy is between *Pseudoalteromonadaceae Vibrionaceae* and *Alteromonadaceae* in the peptide treatment. As mentioned above, all three families were able to grow on peptides in monocultures (Figure 5), but when peptides were added to the EMS natural community *Pseudoalteromonadaceae* became dominant (Figure 4C). Interestingly, two mechanisms by which *Pseudoalteromonas piscicida* strains can kill different *Vibrionaceae* strains (including *Photobacterium*) were recently identified: (1) secretion of antimicrobial substances and (2) direct transfer of apparently lytic vesicles to the surface of competing bacteria, which causes the digestion of cell walls (Richards *et al*., 2017).

## Conclusions

To what extent do heterotrophic bacteria differ in their ability to use the major macromolecular blocks that comprise cell biomass, and that are released when cells die? In the specific community tested here, from the ultra-oligotrophic EMS, amino acids seem to be a major driver of heterotrophic growth. Amino acids induced the highest cell growth and activity (BP), and their own uptake (as BP) was not inhibited by any other molecular class. In contrast, other molecular classes induced less growth and lower activity, and the uptake of glucose was inhibited by other molecular classes. Combining these bulk observations with measurements of 16S rRNA expression suggests that that the overall community responses to different DOM components are due to a complex interplay between changes at the scale of individual cells (e.g. catabolite repression) and shifts in community composition. Assessing the growth of individual strains in the lab on different molecular classes, and genomic analyses, provide only partial explanations to the complex patterns exhibited when natural communities respond to different DOM sources. This highlights the need to explore metabolism across more inter- and intra-clade diversity in marine heterotrophic bacteria, possibly with the help of genome-scale models that can help bridge lab experiments with ‘omics based field measurements.

## Acknowledgements

We thank the captain and crew of the M/V Mediterranean Explorer for help during the cruise, Tal Ben-Ezra and Mike Krom for the nutrient analyses and Yotam Fadida and Yoav Lehahn for the satellite image. This study was supported by the Human Frontiers Science Program (grant number grant RGP0020/2016, to DSh and DSe), the National Science Foundation - United States-Israel Binational Science Foundation (NSFOCE-BSF 1635070 to DSe and Dsh), the Israel Science foundation (grant 1786/20, to DS) and the Southern Marine Science and Engineering Guangdong Laboratory (Guangzhou, grant SMSEGL20SC02, to DS). SG was supported by PhD scholarships from the Yochai Bin Nun foundation (IOLR) and the University of Haifa. The work of EF in Israel was supported by the Association for the Sciences of Limnology and Oceanography (ASLO) LOREX fellowship, NSF-OISE no. 1831075

